# Acute Stress Reduces Reward-Related Neural Activity: Evidence from the Reward Positivity

**DOI:** 10.1101/2020.01.08.898593

**Authors:** Kreshnik Burani, Austin Gallyer, Jon Ryan, Carson Jordan, Thomas Joiner, Greg Hajcak

## Abstract

Stress and blunted reward processing are risk factors for mood disorders, including Major Depressive Disorder (MDD). The experience of acute stress reduces fMRI correlates of reward-related neural activity; however, few studies have examined how acute stress impacts measures of reward derived from event-related potentials (ERPs). The current study examined the impact of an acute stressor on the Reward Positivity (RewP), an ERP that indexes reward sensitivity, in twenty-seven college students. Participants completed a monetary reward task while they placed their left hand in cold water set at 13 degrees Celsius (i.e., acute stress condition) and again while their hand was placed in room temperature water (i.e., control condition). These conditions were separated by one week and performed in a counter-balanced order across participants. The results revealed that the RewP amplitude was blunted in the acute stress condition compared to the control condition. Moreover, there was a trend toward this effect interacting with self-reported depressive symptoms: the RewP was only reduced among individuals who reported low depressive symptoms. The current study suggests that an acute stressor reduces the RewP, and that this effect might be moderated by current depressive symptoms. Future studies might examine the temporal association between reward processing and stress —and how they interact to predict depressive symptoms.

## INTRODUCTION

Stress and aberrant reward system functioning are implicated in the etiology and pathophysiology of Major Depressive Disorder (MDD) (Admon & Pizzagalli, 2015; Hammen, 2005; Whitton, Treadway, & Pizzagalli, 2015). Many studies have found that participants with MDD are characterized by reduced activation of the reward circuit, including blunted activity in the ventral striatum (VS) and the orbitofrontal cortex (Epstein et al., 2006; Forbes et al., 2006; Pizzagalli et al., 2009). Both stress and reward circuit hypoactivation may be risk factors for, and important correlates of, MDD and depressive symptoms. One potential etiological and pathophysiological model of depression that links stress and blunted reward processing to depression postulates that chronic stress reduces reward circuit function and may induce anhedonia (Pizzagalli, 2014)—a core symptom of MDD.

Consistent with this model, previous fMRI research has found that the experience of acute stress reduces neural activity in brain regions involved in reward processing. Lincoln and colleagues (2019) reported blunted dorsal striatum activation to rewards, but not losses, following an acute social stressor compared to baseline (Lincoln et al., 2019). Kumar and colleagues (2014) used a within-subjects design in which participants completed multiple runs of a monetary reward task under stress or no-stress conditions, and reported a trend toward reduced neural activity in the left putamen (a part of the dorsal striatum) during the stress condition compared to the no-stress condition following rewards versus losses (Kumar et al., 2014). Finally, using a physical stressor, Porcelli and colleagues (2012) had adults complete a guessing task where rewards and non-rewards were equiprobable following a stressor (i.e., cold pressor arm wrap maintained at 4 degrees Celsius for 2 minutes) or a control condition (e.g., room temperature arm wrap for two minutes) (Porcelli, Lewis, & Delgado, 2012). In the no-stress group, participants showed greater differentiation between rewards and losses in the orbitofrontal cortex and the dorsal striatum, particularly within the right caudate and the left putamen; on the other hand, the stress group demonstrated a lack of differentiation between rewards and losses within the same brain structures—results that were mainly driven by a decreased response to rewards. Collectively, these studies suggest that acute stress appears to blunt activation of the reward circuit.

Reward circuit function can also be indexed utilizing event-related brain potentials (ERPs). In particular, the reward positivity (RewP) is a positive going ERP that is maximal ~300 ms following the presentation of rewards compared to non-rewards at frontocentral electrode sites (Holroyd, Pakzad-Vaezi, & Krigolson, 2008; Proudfit, 2015). A larger (i.e., more positive) RewP correlates with increased activity in the VS to reward measured using fMRI (Becker, Nitsch, Miltner, & Straube, 2014; Carlson, Foti, Mujica-Parodi, Harmon-Jones, & Hajcak, 2011). A blunted RewP has been linked to personality traits related to depression, such as low positive emotionality and reduced self-reported reward sensitivity (Bowyer et al., 2019; Bress & Hajcak, 2013; Kujawa et al., 2015; Speed et al., 2018; Umemoto & Holroyd, 2017). Indeed, research has also shown that a blunted RewP is evident among clinically depressed children and adults (Belden et al., 2016; Brush, Ehmann, Hajcak, Selby, & Alderman, 2018; Foti & Hajcak, 2009; Liu et al., 2014), is observable prior to the onset of depression, and can predict increases in depressive symptoms and first-onset MDD (Bress, Foti, Kotov, Klein, & Hajcak, 2013; Nelson, Perlman, Klein, Kotov, & Hajcak, 2016). Overall, the above evidence suggests that the RewP reflects neural reward circuit function and is an important individual difference measure of reward sensitivity.

Recently, research has shown that a blunted RewP interacts with increased stressful life events to predict increases in depressive symptoms (Burani et al., 2019; Goldstein et al., 2019). However, the underlying mechanisms of how stress affects reward processing is still relatively unknown—and almost no studies have experimentally examined whether acute stressors affect the RewP. One study by Bodgan et al. (2011) found that the ERP to reward-related feedback was reduced under a threat of shock condition compared to a no-stress condition; however, because only positive feedback was used, it is not clear if this effect was specific to reward-related neural activity (Bogdan, Santesso, Fagerness, Perlis, & Pizzagalli, 2011).

The present study examined the effects of an acute stressor on reward processing, measured using the RewP during a simple guessing task (i.e., the doors task) in which rewards and non-rewards were equally probable. The acute stressor in the current study was a cold pressor which is a robust physical stressor shown to reduce reward-related neural activity using fMRI (Porcelli et al., 2012). Using a within-subjects design, participants completed the doors task during an acute stress condition in which they placed their hand in 13 degrees Celsius water, and again while they placed their hand in room temperature water (i.e., control condition) during a separate testing session separated by one week. We hypothesized that the RewP would be blunted in the stressor condition compared to the control condition. Furthermore, given the documented relationship between increased depression and a reduced RewP, we sought to examine the impact of depressive symptoms on stress-induced reductions in the RewP. Because previous studies have not examined the impact of depression on stress-induced reductions of reward measures, we did not have any *a priori* hypotheses. Because the RewP is blunted in individuals with increased depressive symptoms (Bress and Hajcak, 2013), the effects of the acute stressor might be reduced among individuals with increased depressive symptoms since they are already characterized by decreased reactivity to rewards. Alternatively, increased depression might relate to greater sensitivity to stress (O’Hara, Armeli, Boynton, & Tennen, 2014)—and potentiate the impact of stress on reward-related neural measures.

## METHOD

### Participants

We recruited 33 undergraduate students at a large southeastern university in the United States. Due to missing data (i.e., five participants did not return for a second visit) and noisy EEG data (one participant), we retained 27 participants for all analyses. The final sample consisted of mostly females (61%), and ages ranged from 18 to 20 years (*M* = 18.38, *SD* = 0.5). Exclusion criteria included a history of cardiovascular problems and/or frostbite; however, no participants were excluded for these reasons. The University Institutional Review Board approved all study procedures.

### Measures

#### Patient Health Questionnaire – 8 (PHQ-8) (Kroenke et al., 2009)

The PHQ-8 is an 8-item self-report measure of depression. On the PHQ-8, participants rate how frequently they have experienced each symptom of depression over the previous two weeks, ranging from 0 “*Not at all*” to 3 “*Nearly every day*.” The PHQ-8 has been shown to be a valid and reliable measure that is effective at screening for a diagnosis of depression on par with the longer Patient Health Questionnaire – 9 (Kroenke, Spitzer, & Williams, 2001; Wu et al., 2019). In the present study, the PHQ-8 was administered twice (see Procedure), and demonstrated excellent internal consistency at both time points, (i.e., ω = .88 and ω = .92).

### Procedure

There were two study visits that occurred one week apart. During the visits, participants first completed a brief self-report questionnaire, which included the PHQ-8. Then, participants were connected to a BrainVision LiveAmp EEG system. Next, participants completed a variation of the doors task. This task consisted of seven blocks with six trials in each block. Before each block, participants were instructed to place their left hand in the water, which was set to either 13°C (Acute Stress Condition) or room temperature (Control Condition). On each trial, participants were presented with two doors on the screen and instructed to select one of the two doors using the left or right mouse button. After selecting a door, a fixation cross was presented for 1000 ms. Then, participants were shown either a green arrow pointing up, indicating that they had won $1.00, or a red arrow pointing down, indicating that they had lost $0.50, for 2000 ms. Following feedback, a fixation cross was then presented again for 1500 ms. Between blocks, participants were asked to rate the level of pain they experienced during the previous block on a 0 (least) to 10 (most) Likert scale while their hand was out of the water. The pain rating across blocks was averaged within each condition, and a difference score was calculated (ΔPain Rating). Participants completed the same task at each study visit, with the order of the acute stress and control conditions randomized across subjects.

### EEG Processing and Recording

Electroencephalography (EEG) was continuously recorded while participants completed the doors task. The EEG was recorded using an actiCAP and ten slim electrodes positioned in accordance with the 10/20 system (LiveAmp, Brain Products GmbH, Gilching, Germany). Electrode FCz served as the online recording reference, and a ground electrode was placed on the forehead at FPz. Two electrodes were placed on left (TP9) and right (TP10) mastoids. Electroculogram (EOG) was recorded using four electrodes: two placed approximately 1 cm above and below the left eye and two at the outer canthi of both eyes. The remaining two electrodes were placed on the scalp at Cz and Pz. The online EEG signal was digitized at 500 Hz and band-pass filtered from 0.01 to 100 Hz, while impedances were kept below 25 kΩ.

EEG data were analyzed using BrainVision Analyzer, version 2.1 (Brain Products, Gilching, Germany). The raw data were re-referenced offline to the average of the left and right mastoids and band-pass filtered from 0.1 to 30 Hz. Eyeblink and ocular-movement corrections were performed using established standards described by Gratton, Coles, and Donchin (Gratton, Coles, & Donchin, 1983). Feedback-locked epochs were extracted with a duration of 1000 ms, starting 200 ms before feedback presentation. The 200 ms segment prior to feedback served as the baseline. Epochs containing a voltage greater than 50 μV between sample points, a voltage difference of 300 μV within a segment, or a maximum voltage difference of less than 0.5 μV within 100 ms intervals were automatically rejected. Feedback-locked ERPs were averaged separately for gain and loss trials. For each participant, ERPs to gains and losses were quantified using the average activity between 250 ms and 350 ms following presentation of feedback at FCz. The RewP was calculated as the difference between the ERP to gains minus the ERP to losses.

## RESULTS

### Pain Ratings

As a manipulation check, participants reported experiencing more pain during the acute stress condition (*M* = 4.17, *SD* = 1.98) compared to the control condition (*M* = .20, *SD* = .35; *F* [1, 22] = 110.08, *p* < .001, η^2^ = 0.70)^1^.

### ERP Data

To examine the effects of the acute stress condition on the RewP, we conducted a repeated measures ANCOVA, with condition (acute stress vs. control) entered as within-subjects factor, and the average PHQ-8 score across the two administrations as a covariate. To follow-up a possible interaction between PHQ-8 and condition, we utilized the PHQ-8 score as a moderator. Figure 1 presents ERPs to gains and losses, as well as the gain minus loss difference waveform, during the acute stress and control conditions, respectively. Consistent with the impression from Figure 1B, the RewP (i.e., the difference between gains and losses) was smaller in the acute stress condition (*F* [1, 24] = 4.37, *p* = .047, η^2^ = 0.06). Though it was non-significant, it is interesting to consider the interaction between PHQ-8 and condition (*F* [1, 24] = 2.59, *p* = .121, η^2^ = 0.04). To follow this up, we split the sample using the median PHQ-8 score, and again conducted a one-way repeated measures ANOVA with condition as a within-subjects factor in the high and low depressed groups separately. Figure 2 presents the ERPs to gains and losses in each condition as well as the difference waveform, split by those in the high depressed group and the low depressed group. In the low depressed group, we found a significant main effect of Condition on RewP (*F* [1, 11] = 5.99, *p* = .032, η^2^ = 0.13; see Fig. 2D). In contrast, in the high depressed group, there was no impact of Condition on RewP (*F* [1, 13] < .001, *p* = .984, η^2^ < .01; see Fig. 2B^2^).

**Figure 1:**
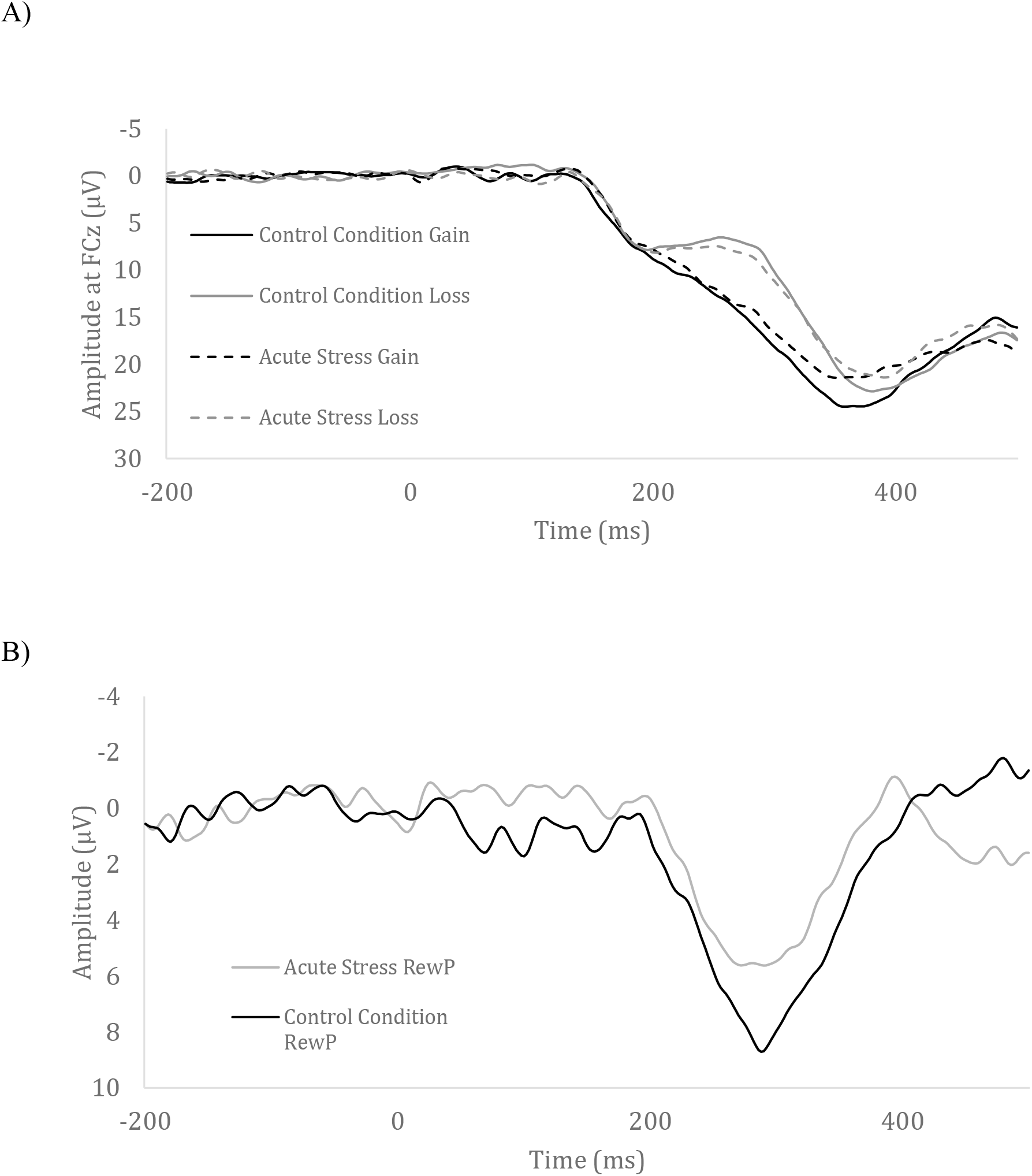
The ERPs to the acute stress win, acute stress loss, control condition win and control condition loss (A). The gain minus loss difference waveform in the acute stress condition and in the control condition (B).

**Figure 2:**
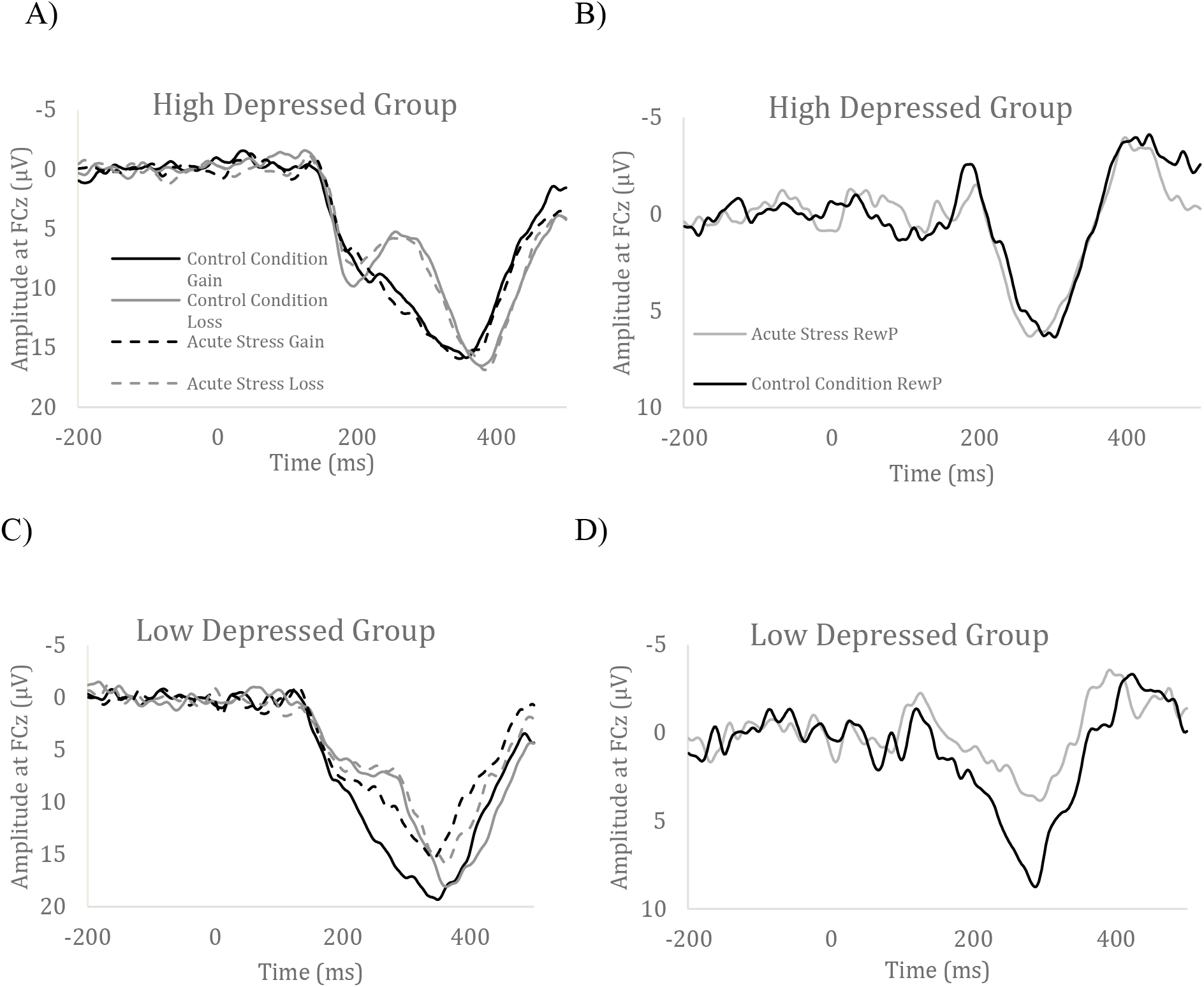
The ERPs to the acute stress gain, acute stress loss, control condition gain and control condition loss in the high depressed group (A) and low depressed group (C) as well as the gain minus loss difference waveform in the high depressed group (B) and low depressed group (D).

## DISCUSSION

The current study examined whether the experience of acute stress impacted reward processing as measured by the RewP. We utilized a within-subjects experimental design where participants completed a reward task during an acute stress (i.e., cold pressor) and a control condition, separated by a week. The results revealed that acute stress reduced the RewP. Since previous studies have found that a potentiated RewP is related to increased activation in the VS (Carlson, Foti, Mujica-Parodi, Harmon-Jones and Hajcak, 2011; Becker, Nitsch, Miltner, and Straube, 2014), the results of the current study provide further support for the notion that reward circuit function is reduced by acute stress. Specifically, the current results are in line with past fMRI research that has shown that acute stress reduces activity in the dorsal striatum and the orbitofrontal cortex following a stressor (Lincoln et al., 2019; Porcelli, Lewis, and Delgado, 2012; Kumar et al., 2014). Furthermore, the present study extends the current literature by demonstrating that ERP measures of reward processing (i.e., RewP) are similarly impacted by acute stress.

Moreover, there was a trend toward depressive symptoms moderating the impact of acute stress on neural measures of reward processing, such that the RewP was blunted in the acute stress condition compared to the control condition in individuals who reported low, but not high, depressive symptoms. A lack of responsivity across the acute stress and control conditions in the high depressed group is consistent with the emotional context insensitivity (ECI) hypothesis of depression, insofar as depressed individuals appear less able to modulate emotional responsivity (Rottenberg, Gross, & Gotlib, 2005). It is also possible that these results could be due to floor effects. A similar and independent effect was evident when considering pain ratings: stress appeared to reduce the RewP only among those who experienced a relatively large change in pain ratings between the acute stress and control conditions. Although these potential moderator effects could shed light on who is most likely to experience stress-induced reductions in reward-related neural activity, these effects require replication given the small sample size for between-subjects associations.

The current study found blunted reward activity in the face of acute stress; however, studies from animal research have demonstrated that reward system function in response to acute stress varies depending on the type of stressor (Pizzagalli, 2014). In response to controllable and short duration stress, dopamine release in the nucleus accumbens is enhanced. This response is thought to be adaptive, insofar as increased dopamine release has been related to active coping strategies and behavioral activation which might aid an organism in escaping or avoiding a stressor (Cabib & Puglisi-Allegra, 2012). Therefore, the effects of acute stress could also be perceived as beneficial – and future work is needed to further examine the how stress-induced reductions in neural reward response relate to the impact of stressors that are more chronic and uncontrollable.

There are several limitations to note in the present study. The current study utilized an unselected college student sample, which could limit the generalizability of our results. Future studies might examine the impact on acute stress on reward processing by comparing individuals with current MDD versus healthy controls to examine the impact of more extreme depressive symptoms on stress-related reductions in the RewP. Because the current study utilized a physical stressor (i.e., cold pressor), it might be important for future studies to compare the impact of other forms of acute stress (i.e., social stress) on reward processing to determine if different stressors result in similar effects. Finally, it is likely that the acute stress in the current study was more distracting than the control condition. Future studies will need to include a distracting, but not stressful, control condition to confirm that distraction itself does not reduce the RewP.

In conclusion, the current study found that acute stress reduces reward processing indexed by the RewP – and that this effect was driven by individuals low in depressive symptoms. These results suggest that increased depressive symptoms might attenuate the impact of acute stress on reward processing. Based on these results, one important question is whether stress-induced reductions in reward processing could predict increases in depressive symptoms over time. In addition, future studies might examine whether stress-induced reductions in the RewP predict stress reactivity to real life stressors. The current study suggests that acute stress induces anhedonia among individuals who report low depressive symptoms – and this may mimic the way in which chronic stressors might blunt reward sensitivity and give rise to symptoms of depression. To further examine this possibility, it will be important to examine the temporal association between stress, reward processing, and changes in depressive symptoms.

## Footnotes

1 Change in pain rating (i.e., acute stress minus control; ΔPain) was uncorrelated with depressive symptom severity (*r* = −.11, *p* = .63).

2 When ΔPain was included as an additional covariate, the interaction between PHQ-8 and condition remained unchanged (*F* [1, 20] = 2.73, *p* = .11). In addition, there was a similar and independent trend toward an interaction between ΔPain and condition (*F* [1, 20] = 2.35, *p* = .14), such that the RewP did not differ between the acute stress and control conditions among those who were below the median on ΔPain ratings (*F* [1, 10] = .43, *p* > .50); however, the RewP was reduced in the acute stress compared to control condition among those above the median on ΔPain ratings (*F* [1, 10] = 3.64, *p* = .09).

## Notes

#### Summary of Updates

Correct author name error.

